# “Immunomodulatory nanoparticles elicit antifibrotic monocyte activation to resolve murine pulmonary fibrosis”

**DOI:** 10.64898/2025.12.22.696047

**Authors:** Hannah Viola, Hannah Carter, Kate Griffin, Rita Medina Costa, Riley McDonald, Brennan Callow, Francina Gonzalez de Los Santos, Marisa Martinez, Ryan Chen, Zharia Hunter, Gary Luker, Bethany B. Moore, Lonnie Shea

## Abstract

Organ fibrosis presents a substantial disease burden with few therapeutic options. Innate immunity mediates fibrinogenesis, but also plays a major role in fibrinolysis. Here, we show that immunomodulatory nanoparticles (NPs) can harness this endogenous antifibrotic capacity by catalyzing monocyte activation leading to resolution of bleomycin-induced pulmonary fibrosis *in vivo*. Cargo-free NPs comprised of the degradable biopolymer poly(lactide-co-glycolide) (PLG) induce a transcriptional shift toward antifibrotic immune activation in profibrotic M2 macrophages (MΦs) *in vitro*. NPs stimulate M2 MΦs toward a glycolytic, rather than fatty acid oxidative, metabolism; suppress canonical M2 markers like arginase-1 (*Arg1*) and periostin (*Postn*); and upregulate collagenases, hyaluronidases and immunoregulatory factors. When delivered intravenously *in vivo*, NPs reverse established bleomycin-induced pulmonary fibrosis and invert the trajectory of over 1,000 genes from pre- to post-treatment according to bulk RNA-sequencing. NPs also suppress profibrotic signaling and increase expression of repair-associated pathways like peroxisome proliferator-activated receptor gamma (PPAR-γ), nuclear retinoic acid receptor (RAR), vascular endothelial growth factor (VEGF), and sphingolipid signaling in fibrotic lungs. Flow cytometry confirms that NPs induce monocyte recruitment to fibrotic lungs *via* enhanced integrin expression. Altogether, NPs induce a robust pro-regenerative signature comprised of ECM degradation, inflammation resolution, and tissue repair pathways, concomitant with increased NP+ monocyte recruitment to fibrotic lungs. This work demonstrates that monocytes are not intrinsically profibrotic, but rather, their effects are context-dependent, and they retain a capacity for fibrotic resolution under conditions that can be induced by materials with translational potential.

**Significance Statement:** Organ fibrosis can follow from tissue injury and substantially impairs organ function, but it is incurable and has limited therapeutic options. Myeloid immune cells drive fibrosis onset and progression, but they can also mediate fibrosis resolution. We used polymeric NPs made from clinically translatable biomaterials to harness the antifibrotic capacity of immune cells as a fibrosis therapy. We found that intravenous NPs can reprogram myeloid cells to acquire an antifibrotic phenotype in a mouse model of pulmonary fibrosis. NPs increased myeloid activation and pulmonary recruitment while decreasing fibrosis, showing that directed immune activation, instead of suppression, could be an effective therapeutic strategy for fibrosis.

## Introduction

Organ fibrosis is a common endpoint of chronic illness that contributes to the disease burden of 1 in 4 individuals worldwide^1^, but has no cure and few therapeutic options. Fibrosis describes tissue scarring that occurs in the context of failed repair after severe or repetitive tissue injury or inflammation. Failed repair drives an inflammatory wound healing response that promotes the deposition of insoluble fibrillar collagens and other extracellular matrix (ECM) components by activated myofibroblasts. This ECM-rich microenvironment is itself profibrotic, perpetuating a cycle of dysregulated repair that ultimately compromises organ structure and function over time^2^. Small molecule inhibition of profibrotic signaling pathways with nintedanib and pirfenidone has slowed progression, but complete resolution remains elusive. The lack of success in treating fibrosis, despite decades of research, may relate, in part, to a limited focus on the promotion of antifibrotic responses. Fibrosis resolution is an active biological process that requires global structural reorganization that is accomplished by cooperation of all tissue cell types toward a homeostatic endpoint, but few therapeutic investigations have aimed to promote this process^2–4^.

Mononuclear phagocytes (MPS) can be targeted to catalyze fibrosis resolution. MPS are central orchestrators of the tissue remodeling process, and as such they play a critical role in both the development and resolution of fibrosis^5^. Indeed, deletion of monocytes or MΦs prior to fibrotic emergence is protective^6^, but their deletion during the spontaneous resolution phase delays resolution and exacerbates fibrosis^7,8^. These observations help explain why several large clinical trials have found little to no benefit for compounds that broadly inhibit inflammatory signaling, such as corticosteroids^9^ and pentraxin-2^10^. These compounds decrease profibrotic inflammation, but similarly prevent antifibrotic immune effector functions that require immune activation, such as tissue recruitment, differentiation, and secretion of inflammation-associated mediators^2,3,11^, yielding little net benefit.

Here, we report that directed activation of myeloid cells with immunomodulatory polymeric NPs is sufficient to ameliorate established pulmonary fibrosis *in vivo* by inciting proresolving immune activation. Polymeric NPs profoundly alter the transcriptome of M2-polarized profibrotic MΦs *in vitro* by modulating their metabolism and signaling architecture to elicit an antifibrotic phenotype consisting of collagenases and immune-resolving mediators. Further, intravenous NPs target peripheral monocytes to induce their recruitment and activation in fibrotic lungs, leading to resolution of pulmonary fibrosis *in vivo*. Bulk RNA sequencing and flow cytometry of NP-treated lungs supports the notion that NPs drive an activated response from monocytes that accompanies fibrosis resolution *via* upregulation of ECM-degrading collagenases like matrix metalloproteinase 13 (MMP-13), collagen-binding phagocytic receptors like LAIR-1, and enzymes that produce immune-resolving lipids like prostaglandin E^2^ (PGE^2^). This work supports the emerging concept that innate immune activation is not universally pathological in the setting of fibrosis, but rather, the type and degree of inflammatory activation determines whether the response is protective or pathologic. These findings motivate renewed enthusiasm for immunomodulatory therapy as a fibrosis treatment, with the goal of harnessing endogenous antifibrotic immunity in a pro-resolving manner rather than broadly suppressing immunity.

## Results

### NPs polarize MΦs toward a proresolving phenotype *in vitro*

We first studied how NPs affect the phenotypic polarization of bone marrow-derived MΦs (BMDMs) *in vitro*. Although BMDM polarization states do not fully recapitulate *in vivo* effects^12^, they can model the effect of NPs on well-characterized myeloid polarization states, which provides insight into their capacity for modulating phenotypes *in vivo*. Namely, we can evaluate how NPs affect BMDM polarization toward stereotypical M1 vs. M2 states that are linked to antifibrotic vs. profibrotic effects *in vivo*, respectively^13–15^. M2-like MΦs have been identified in fibrotic lungs and produce TGF-β, VEGF, arginase, and other profibrotic mediators, whereas M1-like MΦs or those with a “mixed” M1/M2 phenotype can be antifibrotic, producing collagenases, collagen receptors, and immune-resolving mediators^16,17^. These pro- and antifibrotic effects are related to the differences in the M1 vs. M2 states’ metabolic, signaling, and epigenetic architectures. The M1 phenotype, induced by LPS and IFN-γ, utilizes aerobic glycolysis for energy and produces acute-phase response proteins like TNF-α, IL-1β, and reactive oxygen species (ROS)^18^. The M2 phenotype, induced by IL-4 and IL-13, utilizes fatty acid oxidation coupled to mitochondrial respiration for energy, is associated with arginase production and ornithine metabolism, and produces late-stage response mediators with roles in tissue repair, such as growth factors (TGF-α, VEGFs) and inflammation-controlling lipids and cytokines (e.g. specialized proresolving mediators, IL-10)^19^. Thus, we can correlate NPs’ effects to these well-characterized states *in vitro* to determine whether NPs elicit profibrotic vs. antifibrotic phenotypes.

We first studied how NPs modify the phenotype of unpolarized (M0) MΦs toward M1 vs. M2 states. We fabricated polymeric NPs comprised of a poly(lactide-co-glycolide) (PLG) core coated with the surfactant poly(ethylene-alt-maleic anhydride) (PEMA) using oil-in-water emulsion followed by solvent evaporation as previously described^20^ (**Figure 1A**). NPs had an average diameter of 604.5 ± 364.2 nm and surface charge of - 45 ± 12.2 mV (**Figure 1B**). M0 BMDMs were cultured with 200 µg/mL NPs for 24 hours, and their conditioned media was transferred into a myofibroblast scratch wound assay (**Figure 1C**). qRT-PCR analysis of M0 MΦs showed that NPs downregulated the expression of M2 genes *Fizz1* and *Cd206* and upregulated M1 genes *iNos* and *Ccl3*/MIP-1α^21^ (**Figure 1C.*i***). Further, conditioned media from NP-treated BMDMs inhibited the migration of TGF-β-differentiated myofibroblasts in a scratch wound assay, indicating that NPs condition unpolarized BMDMs toward an antifibrotic phenotype that was more M1-like than vehicle-treated MΦs (**Figure 1C.*iii***).

**Figure 1.**
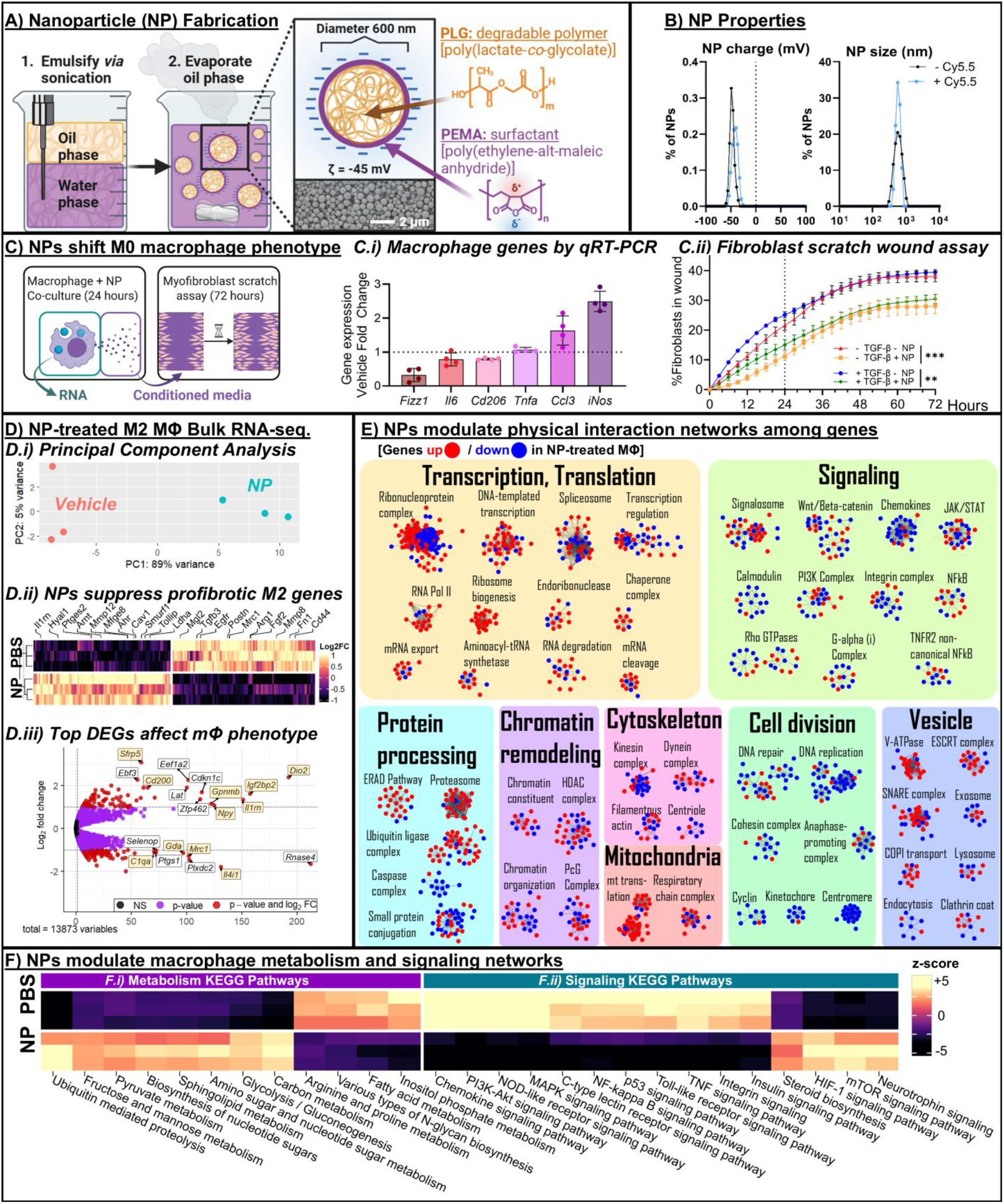
NPs shift M2-polarized macrophages toward fibrosis resolution. **1A)** <u>Nanoparticle fabrication process</u>. Scale bar, 2 µm. **1B)** NP size and surface charge. **1C)** NPs were incubated with M0 BMDMs for 24 hours followed by a myofibroblast scratch wound assay with BMDM conditioned media. **C.*i*** NPs inhibit M2-associated genes by qRT-PCR. **C.*ii*** BMDM conditioned media from NP-treated cells inhibits myofibroblast migration in a scratch wound assay. **1D)** NPs were incubated with M2-polarized BMDMs for 24 hours followed by bulk RNA-sequencing. **D.*i*** <u>PCA of all DEGs</u> finds 89% of variation in first component, demonstrating that most gene expression variation is attributable to NP vs. vehicle conditions. **D.*ii*** <u>Heatmap of all DEGs</u> shows that over 6,000 genes are robustly differentially expressed due to NPs, including upregulated antifibrotic macrophage genes and downregulated classically profibrotic M2 genes. **D.*iii*** <u>Volcano plot of all DEGs</u> emphasizes genes pertinent to macrophage phenotype that are both highly significant (p ≤ 0.05) and ≥1 Log2 fold change in expression. **1E)** <u>Protein-protein interaction network</u> referencing the STRING physical interaction database shows that NPs modulate numerous important cellular functions. NPs upregulate endosome and protein-editing processes, and decrease cell division, signal transduction, and cytoskeletal components. **Statistics: C.*ii*** Two-way ANOVA with post-hoc Tukey’s multiple comparisons test between all groups, α=0.05. *p≤0.05, **p≤0.01, ***p ≤0.001, ****p ≤0.0001. **1A, 1D.i)** Created with Biorender.com under applicable licensure. Data points in **C.*i*** and **D.*i*** represent individual animals.

Next, we investigated the potential for NPs to shift the profibrotic phenotype of M2-polarized MΦs. BMDMs were polarized with IL-4 and IL-13 for 24 hours (10 ng/µL each) and then treated with NPs for 24 hours followed by next-generation bulk RNA sequencing (Illumina). NPs induced 6238 differentially expressed genes (DEGs) (3060 up, 3178 down, **Supplementary File 1**), imparting a substantial transcriptional shift reflected by the robust separation between treatment groups on principal component analysis (**Figure 1D.*i***) and hierarchical clustering of samples on a heatmap including all DEGs (**Figure 1D.*ii***).

NP-induced DEGs reflect a shift in M2 MΦs toward a fibrosis-resolving phenotype with mixed M1 and M2 characteristics that comprises ECM-degrading enzymes and inflammation-resolving mediators (**Table S1**). NP-treated M2 MΦs upregulated ECM-degrading cathepsins, matrix metalloproteinases (MMPs), and hyaluronidases (e.g. *Ctsk, Mmp12, Hyal1)*^16,22^, and the surface receptor *Mfge8* that binds collagen for phagocytotic clearance^23^. Concomitantly, NPs downregulated expression of ECM components including collagen, fibronectin, and chondroitin sulfate proteoglycan (e.g., *Col14a1, Fn1, Cspg5*). NPs also upregulated immunomodulatory receptors and ligands that have antifibrotic roles, such as the canonical aryl hydrocarbon receptor pathway^24^, the enzyme *Ptges2* that produces prostaglandin E^2^, and *Tollip*, a negative regulator of toll-like receptor (TLR) signaling whose dysfunction is one of only a few known genetic risk factors for developing pulmonary fibrosis^25–27^. Likewise, NPs downregulated profibrotic mediators including the growth factors TGF-β and periostin, numerous CCL- and CXCL-chemoattractants like *Ccl2* and *Ccl12*, and *C1q* complement components^28,29^ (**Table S1**). NPs also reduced stereotypical M2-associated genes *Arg1, Retnla*/FIZZ-1, and *Cd163*. However, NPs also downregulated M1-associated genes, like *Il1b, Stat1*, and *Jak2*, suggesting that NPs do not uniliaterally increase M1 phenotypic features as a consequence of decreasing M2 markers. However, a volcano plot shows that genes with the largest fold-change and lowest p-value are related to pro- or anti-fibrotic effects mediated by MΦs (**Figure 1D.*iii***). Namely, NPs upregulate antagonists for profibrotic Wnt/β-catenin (*Sfrp5)* and IL-1 (*Il1rn, Npy*)^30–33^; inflammation controlling mediators (*Gpnmb*^34,35^, *Cd200*^36^, and *Dio2*^37,38^); and the HIF-1α-stabilizing factor *Igf2bp2* that promotes glycolytic metabolism^39,40^. In contrast, the volcano plot highlights the downregulation of *Mrc1*, a prototypical M2 marker gene^41^; *Il4i1*, which promotes M2 marker expression^42^; *Gda*, which promotes profibrotic IL-6 production^43^; and *C1qa*, which activates fibroblasts and is elevated in pulmonary fibrosis patients^28,29,44^. Taken together, these results suggest that NPs remodel MΦ phenotype from a profibrotic M2 state to a matrix-degrading, inflammation-resolving state that does not clearly align with either M1 or M2 phenotypes.

Accompanying these changes, we observed remodeling of genes controlling transcriptional regulation. NPs upregulated the transcription factor (TF) peroxisome proliferator-activated receptor gamma (PPAR-γ), which is central to MΦs’ roles in resolving inflammation, fibrosis, and wound repair^16,45–47^, and shifted expression of TF lineages that orchestrate inflammatory responses and cell differentiation trajectories (i.e., Janus-associated kinase (JAK), JAGGED (JAG), signal transducer and activator of transcription (STAT), CCAAT/enhancer-binding proteins (CEBP), and interferon regulatory factors (IRF)) (**Table S1**). NP-treated MΦs also downregulated histone deacetylases, which have notably been a target of antifibrotic therapy^48^, and increased expression of lysine demethylases (*Kdm*s) and histone acetyltransferases (*Kat*s), suggesting NPs stimulate remodeling of the epigenome. Finally, NPs increase DEAD box helicases that modify RNA transcription and metabolism to regulate innate immunity^49^. These changes reflect reworking of transcriptional architecture that accompanies substantial shifts in cellular function.

NPs also modified signaling architecture, especially tied to metabolism. We performed GSEA referencing the Kyoto Encyclopedia of Genes and Genomes (KEGG) to identify significantly enriched pathways in NP-vs. vehicle-treated M2 MΦs. NPs terms related to glycolysis, pyruvate metabolism, and nucleotide sugar metabolism and biosynthesis (**Table S2, Supplemental File 2**), and downregulated pathways related to fatty acid oxidation, arginine/proline metabolism, and oxidative phosphorylation. These metabolic shifts are consistent with NP-mediated upregulation of M1-associated pathways, especially the concomitant upregulation of NFκB, HIF-1α, and mTOR cascades. NPs also downregulated M2-associated VEGF, MAPK, TLR, and PI3K/AKT cascades. However, NPs also upregulated the M2-promoting p53 pathway and downregulated M1-associated TNF signaling, showing that the polarization imparted by NPs is not unilaterally towards an M1-like phenotype. To further characterize changes in cellular architecture, we constructed a molecular interaction network between all DEGs referencing the STRING database of curated protein-protein binding relationships^50,51^, identified non-interacting clusters of DEGs with similar functions based on gene set enrichment analysis (GSEA), and manually assigned clusters to major categories of cellular function. NPs suppressed genes related to cell division and signal transduction, and enhanced genes related to gene expression, protein editing, vesicle processing, and mitochondria (**Figure 1E**). These changes reflect a shift in cellular activity away from cell division toward vesicle-mediated transport (e.g., SNARE complex, ESCRT complex, COPI transport), increased protein turnover (ribosome biogenesis, mRNA export, proteasome, ubiquitin ligase complex), and activation of mitochondria (mt translation).

Taken together, these findings show that the NPs substantially modulate transcriptional signatures of MΦ phenotype and effector functions by influencing metabolism, signaling, and epigenetic wiring. These changes promote a matrix-degrading, inflammation-resolving phenotype that accompanies an increase in M1-like features like glycolytic metabolism and a decrease in stereotypical M2 features like FIZZ-1, arginase, and fatty acid oxidative metabolism. However, NPs do not clearly induce an M1 phenotype, but rather produce a mixed phenotype. This is consistent with reports that antifibrotic macrophages adopt a mixed M1/M2 phenotype accompanied by ECM degrading enzymes and collagen binding receptors^16^.

### Intravenous immunomodulatory NPs resolve *in vivo* pulmonary fibrosis at a late intervention stage

Since NPs modify MΦs *in vitro*, we investigated the capacity for NPs to exert antifibrotic effects on peripheral monocytes in fibrotic animals. Peripheral monocytes migrate into the fibrotic lung, where they become profibrotic monocyte-derived MΦs with an M2-like phenotype^14,52^. Based on our *in vitro* results, we hypothesized that targeting of peripheral myeloid cells with NPs during or prior to the fibrotic phase of bleomycin-induced pulmonary fibrosis would promote resolution by shifting myeloid phenotypes and trafficking patterns to both reduce their pathogenicity and promote immune-mediated resolution. NPs were fabricated with Cy5.5-labeled PLG for *in vivo* tracking, which did not affect NP properties (**Figure 1B**). Biodistribution studies confirmed *in vivo* degradation of NPs from the lungs, liver, spleen, and kidneys with half lives ranging from ∼1-3 days (**Figure S1**).

The efficacy of intravenously delivered NPs at preventing and reversing pulmonary fibrosis was evaluated using early and late dosing strategies, respectively (Figure 2C)^53^. Pulmonary fibrosis was induced using the oropharyngeal bleomycin model as previously described (0.0125 U/mouse ∼ 0.5 U/kg)^54^. NPs were delivered intravenously via retro-orbital injection at 2 mg/dose in phosphate-buffered saline (Vehicle) beginning on day 7 (early) or day 14 (late), followed by endpoint analysis on day 21 post-bleomycin (**Figure 2A**). Both early and late NPs reduced collagen deposition by Day 21 as quantified with whole lung hydroxyproline assay (**Figure 2B**). Subsequent analyses were conducted using the late dosing strategy (days 14-18) which corresponds to established fibrosis^53^. Lungs treated at the late timepoint trended towards improved lung ventilation and reduced ventilation heterogeneity on Day 29 post-bleomycin according to μCT imaging and ventilatory perfusion analysis (Figure 1D)^55^. We also found that NPs do not affect clearance of *S. aureus in vivo* in fibrotic mice (**Figure S2**). Together these findings demonstrate that negatively charged, degradable polymeric NPs without therapeutic cargo can interrupt pulmonary fibrosis to reverse lung collagen deposition when administered systemically at a therapeutic timepoint without compromising host immunity.

**Figure 2.**
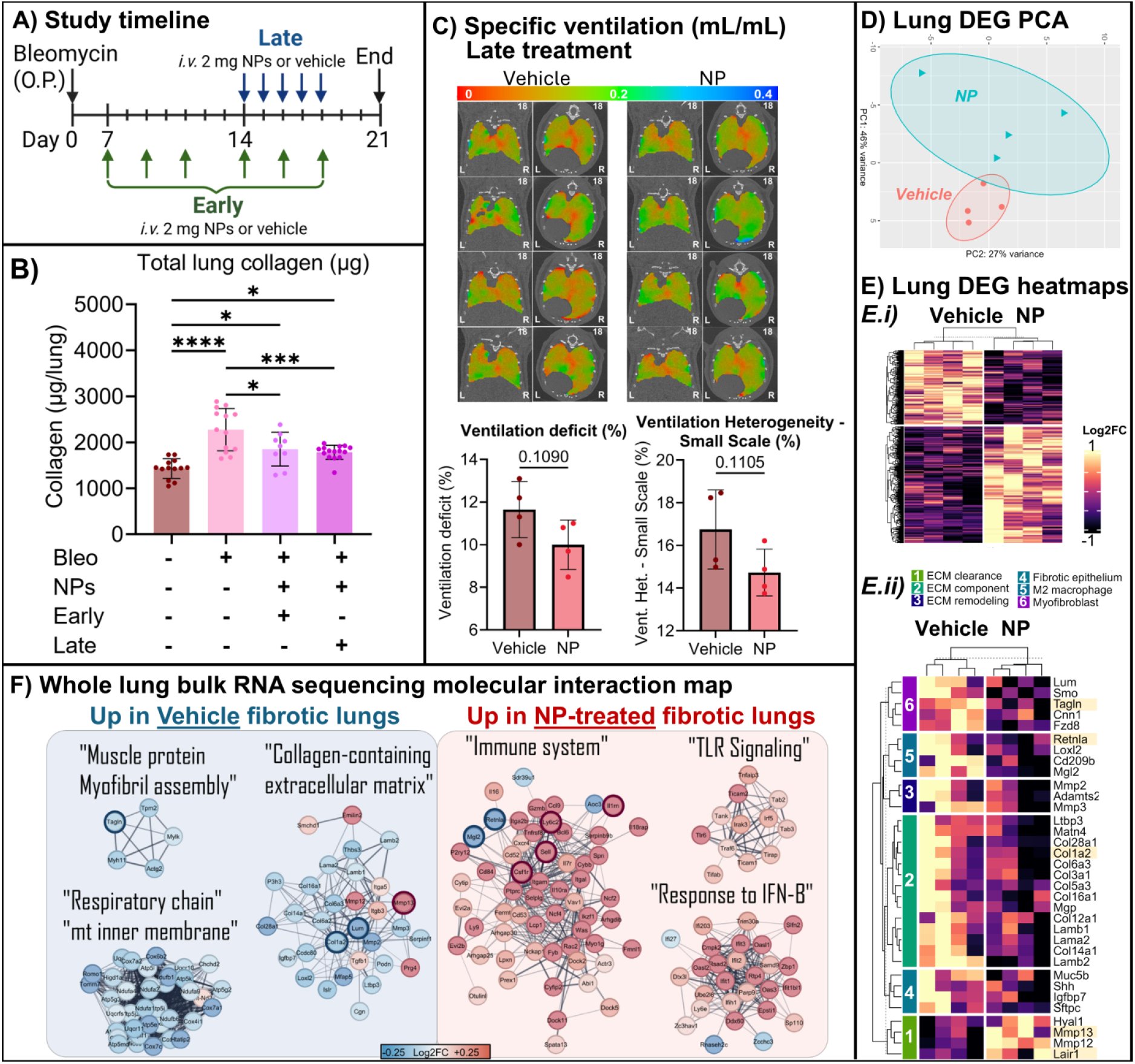
NPs attenuate pulmonary fibrosis *in vivo* at a therapeutic timepoint and induce antifibrotic immune activation. **2A)** <u>Study timeline</u> showing early vs. late delivery of NPs. **2B)** <u>Whole lung collagen</u> measurement by hydroxyproline assay demonstrates that NPs at either timepoint ameliorate fibrosis. **2C)** <u>4D µCT ventilation analysis</u> demonstrates a trend toward improved ventilatory parameters in NP-treated animals. **2D)** PCA Plot of all DEGs shows separation between NP- and Vehicle-treated lungs. Solid ovals, 80% confidence interval (CI); dotted ovals, 95% CI. **2E.*i)*** <u>Heatmap of all lung DEGs</u> shows treatment group separation using unsupervised k-means clustering. Scale, -1 to 1 normalized Log2FoldChange. **2E.ii)** Heatmap shows downregulation of fibrosis-associated genes (*Col1a2, Retnla, Tagln*) and upregulation of innate immune genes as well as immune-associated collagen-sensing (*Lair1*) and collagenase (*Mmp13*) genes. Scale, -1 to 1 normalized Log2FoldChange. **2F)** <u>Whole lung bulk RNA-seq protein-protein interaction analysis</u> finds downregulation of networks for collagen production program, muscle fiber machinery, and mitochondrial respiration, and upregulation of networks for innate immune activation. Statistics: B) One-way ANOVA with post-hoc Tukey’s multiple comparisons test between all groups, α=0.05. *C)* Student’s t-test, α=0.05. *p≤0.05, **p≤0.01, ***p ≤0.001, ****p ≤0.0001. **2A)** Created with Biorender.com under applicable licensure. Data points in **B)** and **D)** represent individual animals.

### Trajectory analysis reveals NP-driven resolving immune activation signature

We next assessed how NP delivery drove the pulmonary transcriptional landscape towards fibrosis resolution by conducting bulk RNA-sequencing on whole lung homogenates from 3 conditions: Day 14 (pre-treatment, n=4), Day 21 NP (n=4), and Day 21 Vehicle (n=4) (mean 44 ± 7.7 million reads/sample). First, we compared NP-vs. Vehicle-treated lungs on day 21 and identified 1403 DEGs (862 up, 541 down, **Supplementary File 3**) *via* Likelihood Ratio test in DESeq2^56^ with independent hypothesis weighting^57^. A PCA plot of these DEGs shows distinct transcription between groups (**Figure 2D**), and unsupervised k-means clustering separates treatment groups on a heatmap of all DEGs (**Figure 2E.*i***). A second heatmap (**Figure 2E.*ii***) highlights the downregulation of genes associated with myofibroblasts, epithelial-mesenchymal transition, and M2 MΦs (**Figure 2D, Table S3**) and shows that NPs increase transcripts for matrix-degrading enzymes like the collagenase *Mmp13* and collagen receptor *Lair1* (**Figure 2C, Table S3**). To understand signaling networks in NP-treated lungs, we constructed a protein-protein interaction network referencing the STRING database and characterized clusters using GSEA. This network showed that NPs downregulated transcriptional networks related to myofibrils, collagen-containing ECM, and mitochondrial respiration, and upregulated a large network of related immune signaling pathways (**Figure 2F**). NPs also upregulated a TLR signaling network and a cluster of interferon (IFN) stimulated genes that GSEA labeled “response to IFN-β”. Type I interferons, including IFN-β, have antifibrotic effects *in vivo*^58,59^. These data suggest that in murine lungs, NPs decreased profibrotic signaling and ECM deposition and increased ECM-degrading immunoregulatory pathways, similar to our findings *in vitro* in M2 polarized MΦs.

We next investigated the changes in NP-treated lungs pre-vs. post-treatment to understand how NPs alter the trajectory of gene expression in fibrotic lungs. Between NP and vehicle-treated lungs on Day 21, the majority of DEGs (60.4%) and enriched pathways (86.6%) were upregulated in NP-treated lungs, suggesting that NPs stimulate an active transcriptional response that reshapes the pulmonary environment, as opposed to passively inhibiting fibrogenic processes. Consensus clustering of day 21 DEGs was performed using the R package DEGReport^60^ to characterize the trajectory of DEGs from day 14 to day 21 (**Figure 3A-C**). Most DEGs induced by NPs represent a departure from the Day 14 profibrotic milieu. Clusters 1 and 3, which comprised the majority of DEGs, captured genes that were increased (673 genes) or decreased (477 genes) by NPs but remained relatively unchanged in vehicle-treated lungs from Day 14 to Day 21 (**Figure 3B,C**). Clusters 2 (189 genes) and 4 (64 genes) captured genes preserved near Day 14 levels in NP-treated, but not vehicle, lungs. This analysis demonstrated that NPs primarily affect transcription by inducing novel gene expression programs that are absent in the untreated lungs at both timepoints, effectively pushing the lungs further away from pre-treatment conditions by either up- or down-regulating over 1,000 genes that would otherwise maintain similar or lower expression by day 21.

**Figure 3.**
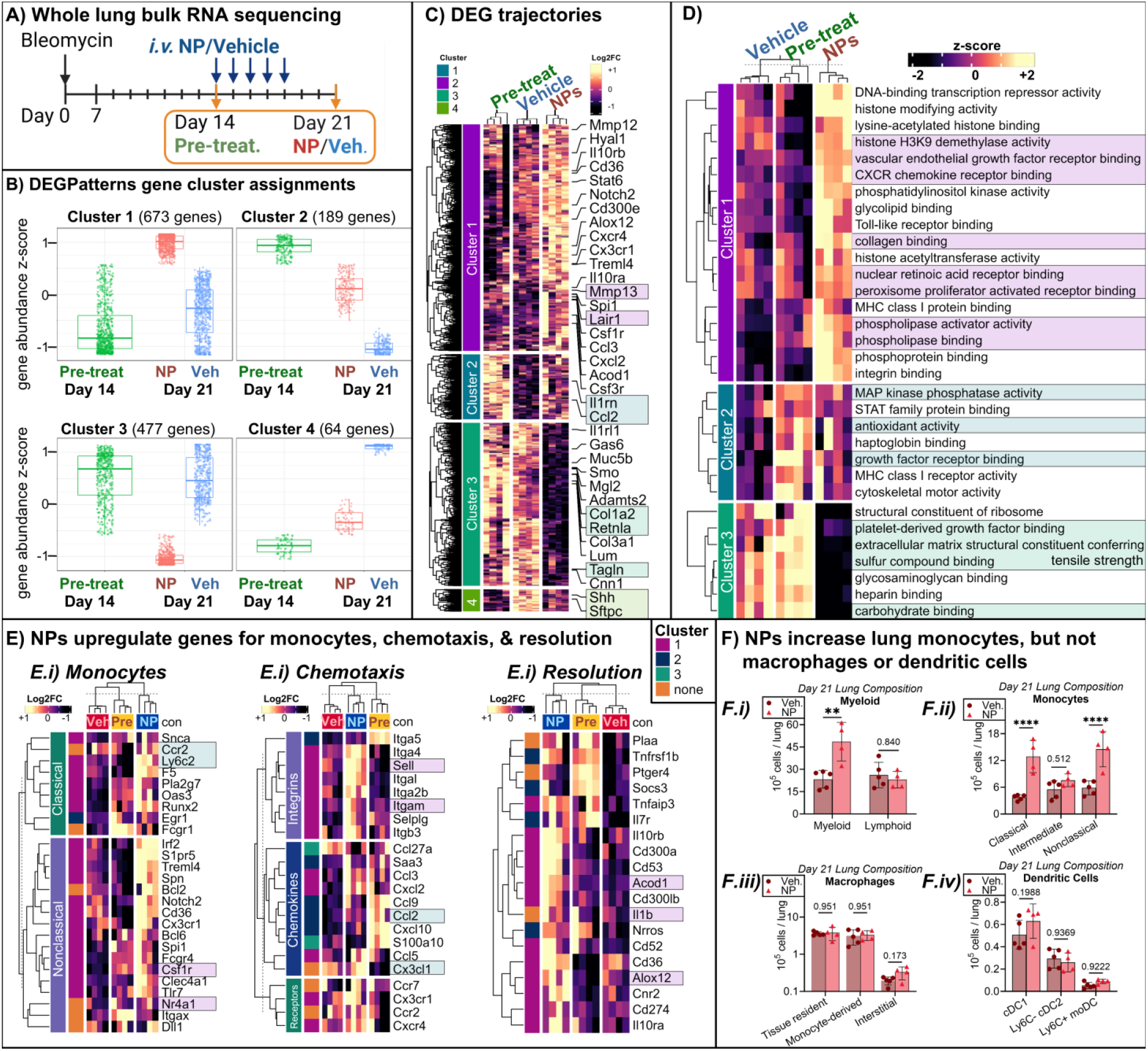
NPs alter fibrotic trajectory towards inflammation resolution. 3A) <u>Study timeline</u> showing sequencing timepoints. **3B)** <u>DEG trajectory analysis</u> shows that NPs change the trajectory of over 1,000 genes in Clusters 1 and 3. **3C)** <u>DEG Heatmap with Day 14</u> demonstrates that the majority of genes with reversed trajectories concern immune-mediated fibrinolysis (cluster 1) and fibrosis (cluster 3). **3D)** <u>Heatmap of enriched pathways by trajectory cluster</u> shows that NPs stimulate proresolving mediators like PPAR pathway, retinoic acid, and phospholipase activation in addition to collagen binding and epigenetic editing. NPs preserve early MAPK inhibition, antioxidant and regenerative responses, but suppress fibrotic ECM deposition and PDGF signaling. **3E)** NP-treated lungs exhibited signatures for monocyte recruitment and antifibrotic resolving actions by myeloid cells. **3F)** Flow cytometry confirmed that NP-treated lungs exhibit increased numbers of lung myeloid cells and monocytes, but not macrophages or dendritic cells. **Statistics: *3F*.*i, iii, iv*)** Two-way ANOVA with post-hoc Tukey’s multiple comparisons test between conditions within each subset, α=0.05. ***3F*.*iv)*** Student’s t-test, α=0.05. *p≤0.05, **p≤0.01, ***p ≤0.001, ****p ≤0.0001. **3A)** Created with Biorender.com under applicable licensure. Data points in **F)** represent individual animals.

Signaling networks in these trajectory clusters revealed that NPs activate a proresolving tissue remodeling process promoting tissue homeostasis. Genes in Cluster 1 included the leukocyte-specific collagen-binding receptor *Lair1*^61^; the collagenases MMP-12 and MMP-13; hyaluronidase *Hyal1*; and leukocyte-specific gelatinase MMP-25/leukolysin^62^. Cluster 1 also contained inflammation-resolving mediators like *Alox12*^63^, *Cybb*/*Nox2*^64^, *Ccdc88a*/Girdin^65^, and *Acod1*/*Irg1*^66,67^ (Table S3). Enriched KEGG pathways for Cluster 1, upregulated in NP-treated but not vehicle or pre-treatment lungs, included collagen binding, retinoic acid receptor, phospholipase, peroxisome proliferator-activated receptor (PPAR), sphingolipid, and VEGF pathways (**Figure 3C-D, Table S4, Supplementary File 4**), consistent with lung injury resolution programs^68,69^. Other enriched pathways in Cluster 1 related to innate immune activation including leukocyte adhesion, recruitment, efferocytosis, and receptor-mediated phagocytosis (**Table S4**). Cluster 1 was also enriched in expression of innate immune sensors (NOD-like receptor, Toll-like receptors, C-type lectin receptors), and signaling cascades (JAK-STAT, NFκB) (**Table S4**). Cluster 2, preserved from Day 14 in NP-but not vehicle-treated lungs, included antioxidant activity, growth factor receptor binding, and TNF signaling. Cluster 3 (477 genes) was significantly downregulated from day 14 to day 21 in NP but not vehicle lungs and reflected pathological fibrotic processes. Enriched pathways were related to PDGF, ECM production, and sensing of ECM components like glycosaminoglycans, carbohydrates, heparin, and sulfur compounds (**Figure 3C-D, Table S4**). Cluster 4 was only enriched for calmodulin binding GO pathways. Together with decreased total lung collagen and lower expression of fibrosis-associated genes, these transcriptional analyses are consistent with our *in vitro* findings that suggested NPs activate an antifibrotic innate immune activating response that rewires tissue-level function toward resolution of pulmonary fibrosis.

### NPs stimulate lung recruitment of classical and nonclassical monocytes

Bulk RNA sequencing indicated antifibrotic innate immune activation reminiscent of our *in vitro* findings in M2 MΦ, which motivated studies on monocyte recruitment and activation in fibrotic lungs *in vivo*. We found that NPs induced a transcriptional signature for monocyte recruitment in fibrotic lungs, as well as signatures indicating elevated chemotaxis and secretion of proresolving mediators (**Figure 3E**). Flow cytometry of lungs on Day 21 confirmed that total myeloid cell number in NP-treated lungs was increased (Gating strategy, **Figures S4-S5**). Among myeloid cells, classical and nonclassical monocytes were increased by NPs (**Figure 3F.*i-ii***), while the number of DCs, MΦs, and granulocytes were unchanged (**Figure 3F.iii-iv, Figure S3**). Total lymphoid cell numbers were also unchanged by NPs (**Figure 3F.i**). An increase in monocytes in NP-treated lungs is consistent with elevated expression of myeloid chemoattractants and integrins in NP-vs. vehicle-treated lung RNA sequencing data (**Figure 3E.*i***). Taken together, NPs increase total lung monocyte recruitment to fibrotic lungs as profibrotic transcription decreases.

To identify whether monocytes were directly associated with NPs, we performed flow cytometry on lungs after treatment with fluorescently tagged NPs on day 14 (1 dose) or days 14-18 (5 NP doses) (**Figure 4A**). We identified major myeloid subsets in the lung using established markers (**Tables S5, S6**; **Figures S4-S6**)^70–72^. The majority of recruited NP+ lung cells on day 15 and day 21 were nonclassical monocytes (NCMOs), but not classical monocytes, followed by neutrophils and dendritic cells (**Figure 4B-C**). NPs were sparsely found in MΦ subsets despite significantly increased monocyte numbers and high %NP+ in nonclassical monocytes (**Figure 4B-C**), suggesting that NP+ monocytes are unlikely to become profibrotic monocyte-derived alveolar macrophages (moAMs)^52^.

**Figure 4.**
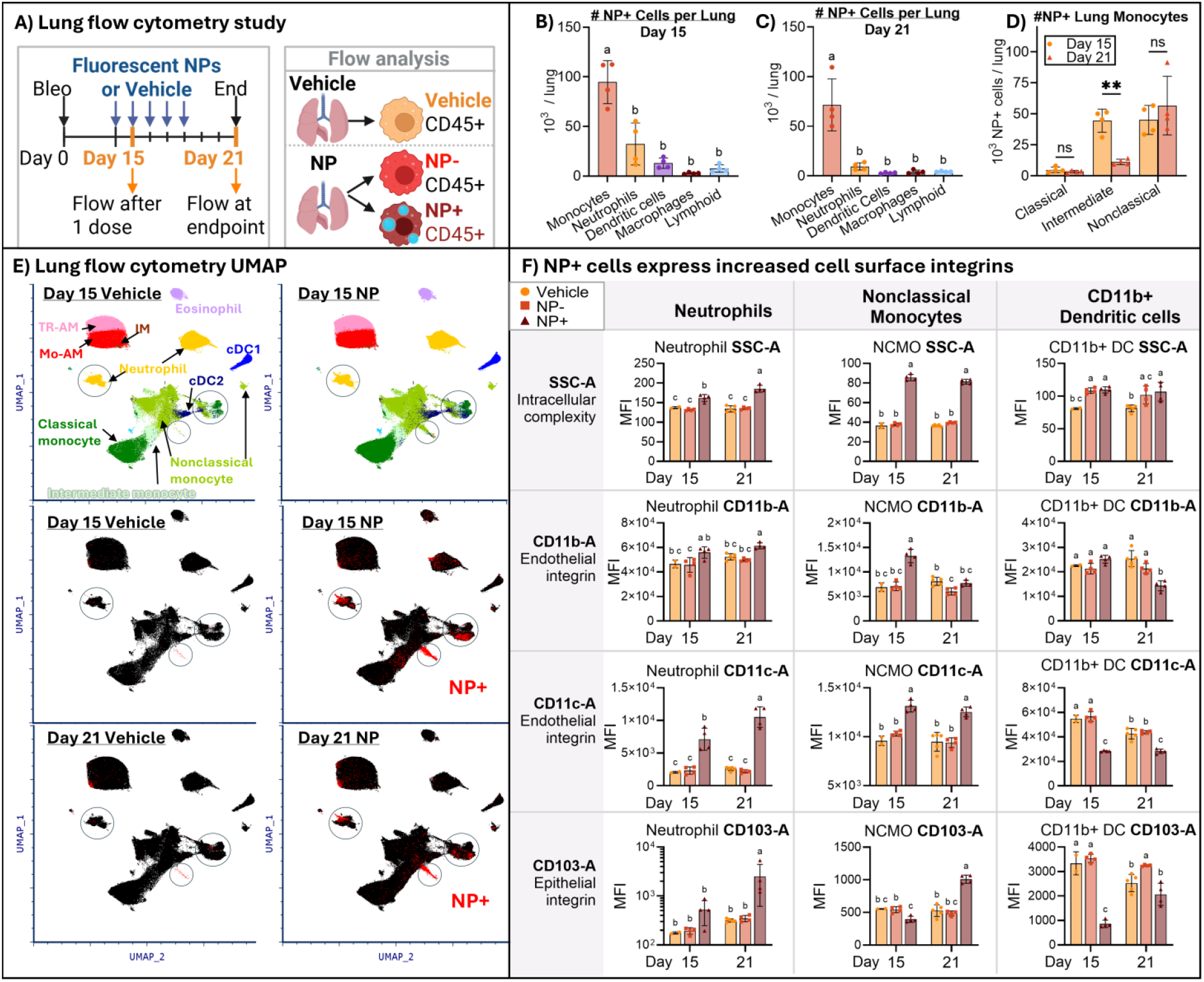
NPs reside primarily in recruited monocytes. 4A) <u>Study timeline</u> showing flow cytometry timepoints. **4B, C)** <u>Composition of NP+ lung cells on Day 15 (</u>**<u>4B</u>**<u>) and Day 21 (</u>**<u>4C</u>**) shows that NPs are found mostly in lung-recruited monocytes. **4D)** NP+ lung monocytes are mostly intermediate or nonclassical. By Day 21, most NP+ cells are nonclassical monocytes. **4E)** <u>UMAP of lung myeloid cells</u> from flow cytometry data was constructed while excluding NP MFIs. UMAP segregates major cell types and identifies that NP+ cells form phenotypically distinct projections in monocytes and neutrophils at both timepoints. **4F)** <u>Surface expression of integrins</u> is upregulated by NP+ lung myeloid cells (NP+), compared to both NP-cells in treated lungs (NP-), and all cells in untreated lungs (Vehicle). **Statistics: 4B**,**C)** One-way ANOVA with post-hoc Tukey’s multiple comparisons test between all groups, α=0.05. **4D**,**F)** Two-way ANOVA with post-hoc Tukey’s multiple comparisons test between conditions, α=0.05. Letters represent statistically significant groups. *p≤0.05, **p≤0.01, ***p ≤0.001, ****p ≤0.0001. **4A)** Created with Biorender.com under applicable licensure. Data points in **B-D)** and **F)** represent individual animals.

The surface expression by myeloid cells from NP-treated lungs was assessed to determine whether NPs directly influenced cell phenotypes upon reaching the lung. We analyzed neutrophils, NCMOs, interstitial monocytes (INTMOs), and monocyte-derived dendritic cells (moDCs), as these subsets had appreciable NP+ cells in the lungs. Uniform manifold approximation and projection (UMAP) of CD45+ myeloid cells revealed that NP+ lung-recruited monocytes and neutrophils are phenotypically distinct from NP-cells in the same lungs. The UMAP was constructed based on scaled, normalized mean fluorescence intensity (MFI) of all surface markers plus side scatter and forward scatter, excluding NP-Cy5.5 MFI. This analysis revealed that NP+ lung recruited monocytes and neutrophils formed projections that extended to non-overlapping regions in the UMAP, showing that they deviate in MFI and/or scatter properties from NP-cells. Analysis of neutrophils, monocytes, and DCs from the circled regions of the UMAP (**Figure 4E**) revealed that NP+ monocytes and neutrophils (“NP+” on **Figure 4F** legend) have greater side scatter and integrin expression than either NP-cells from the same lungs (“NP-”) or vehicle treated lungs (“Veh”) (**Figure 4F**). CD11b+ DCs that were NP+ did not differ substantially from NP- or vehicle lungs, consistent with their greater overlap on UMAP with NP-DCs.

## Discussion

Organ fibrosis is a prevalent endpoint of chronic illness worldwide and has few therapeutic options with limited efficacy that delay, but do not resolve, progression of fibrosis. Tissue-recruited monocytes drive fibrogenesis, but inhibition of innate signaling with broad-spectrum immunomodulators like corticosteroids and pentraxin-2 did not ameliorate fibrosis in clinical trials. These results support a critical role of monocytes in the resolution of fibrosis, which has been demonstrated in multiple animal models^73^. Despite the importance of innate immune activation in mediating fibrosis resolution, few studies have aimed to enhance this endogenous antifibrotic function as a therapeutic strategy.

In this study, we report that pharmacologic-free NPs comprised of the biomaterial PLG have immunomodulatory effects that led to attenuated lung collagen deposition and a trend toward improved lung ventilation. RNA sequencing of NP-treated M2 MΦs showed that NPs reprogram metabolism, signaling networks, and epigenetics to favor an ECM-degrading, inflammation resolving phenotype with suppressed M2 MΦ markers (**Figure 1**). *In vivo*, whole lung RNA sequencing showed that NPs recapitulate their *in vitro* effects by reducing M2 MΦ markers, ECM expression, and fatty acid oxidative metabolism while increasing proresolving immunity and ECM-degrading enzymes, leading to reduced total lung collagen on Day 21 (**Figure 2**). Transcriptional comparison of lungs from pre-to post-treatment showed that NPs uniquely stimulate this innate immune response rather than preserving existing antifibrotic responses (**Figure 3**). Finally, flow cytometry found that total lung monocyte numbers are increased in NP-treated animals, and most NP+ lung cells are NCMOs. MFI analysis found that NP+ myeloid cells increase expression of integrins like CD11b, CD11c, and CD103 (**Figure 4**). Taken together, we conclude that NP treatment after the establishment of ongoing pulmonary fibrosis on day 14 post-bleomycin induces peripheral monocyte activation and recruitment into fibrotic lungs to mediate resolution of the fibrotic milieu and a reduction in total lung collagen.

NP treatment doubles the number of classical monocytes (CMOs) in fibrotic lungs while exerting antifibrotic effects (**Figure 3F**). This result is unexpected based on prior findings that classical monocytes and their derivative macrophages (moAMs) drive fibrogenesis, and ablation of these populations prevents fibrosis^6,7,52,70^. Our results could be explained by the fact that, despite increased CMOs, NP-treated lungs did not have increased moAMs or moDCs. Instead, NP-treated lungs had twice as many NCMOs, which also derive from CMOs, and are considered inflammation-resolving and tissue regenerative^74^. In liver and lung fibrosis, Ly6C^lo^ monocytes/macrophages similar to NCMOs have been identified as mediators of spontaneous resolution^7,75–77^. Therefore, the NP-mediated increase in total lung NCMOs could contribute to NPs’ effects in mediating fibrinolysis.

We showed that NPs bias MPS toward an ECM-degrading, immunoregulatory state in both M2 MΦs *in vitro* and lung-recruited NCMOs *in vivo*, and stimulated M0 MΦs to inhibit fibroblast migration, despite appreciable biological differences between these cell states. Therefore, NPs’ effects are relatively independent of the initial MPS state, suggesting that NPs stimulate a strong, specific response program that triggers antifibrotic and immunoregulatory effects relatively decoupled from the state of MPS prior to NPs. It is likely that such a broad effect is triggered by NPs activating conserved cellular pathways that are universally triggered in MPS in response to specific stimuli. We showed that NPs upregulate the pleiotropic immunoregulatory transcription factor PPAR-γ, a master regulator of tissue-regenerative programming in myeloid cells that is triggered endogenously by phagocytosis of apoptotic cell debris. Additionally, post-phagocytic ‘satiated’ MΦs have fibrinolytic properties^75,78,79^. NPs have comparable size and surface charge as apoptotic bodies, raising the possibility that NPs trigger efferocytosis-like programming in MPS. This would be consistent with increased expression of ECM degradation and inflammation-resolving patterns that we observed across multiple MPS *in vitro* and *in vivo*.

In summary, we showed that NPs comprised of degradable biomaterials can activate monocytes to resolve established pulmonary fibrosis *in vivo*. Therapeutic promotion of endogenous antifibrotic immunity holds potential to complement antifibrotic therapy to facilitate wholistic resolution of pulmonary fibrosis. The NPs are comprised of PLG, an FDA-approved biomaterialabiomaterial that has already been used in human NP-based therapies.

The profound immunomodulatory effects described here underscores the opportunity for cargo-free PLG NPs to be a safe and effective regenerative therapeutic for tissue fibrosis.

## Author CRediT Statement

**Hannah Viola:** Conceptualization, Methodology, Software, Validation, Formal analysis, Investigation, Data Curation, Writing – Original Draft, Writing – Review & Editing, Visualization, Project Administration. **Hannah Carter:** Methodology, Investigation. **Kate Griffin:** Investigation. **Rita Medina Costa:** Investigation, Formal Analysis. **Riley McDonald:** Investigation, Formal Analysis. **Brennan Callow:** Investigation. **Francina Gonzalez de Los Santos:** Methodology, Investigation. **Marisa Martinez:** Investigation. **Ryan Chen:** Investigation. **Zharia Hunter:** Investigation. **Gary Luker:** Supervision, Resources. **Bethany Moore:** Conceptualization, Methodology, Resources, Writing – Review & Editing, Supervision, Funding acquisition. **Lonnie Shea:** Conceptualization, Methodology, Resources, Writing – Review & Editing, Supervision, Funding acquisition. All authors approved the final manuscript.

## Acknowledgements

Bulk RNA library preparation and next-generation sequencing were carried out by the Advanced Genomics Core at the University of Michigan. Flow cytometry data were collected within the University of Michigan’s Flow Cytometry Core Facility. Figures were created with Biorender.com under applicable licensing agreements. The authors acknowledge funding from The University of Michigan Multidisciplinary Training in Lung Disease (5T32HL007749-33) to HV, R35HL176572 to BBM, and R01EB036030 and R01AI148076 to LS.

## Declaration of Interests

Author L.D.S. consults and has financial interests in Cour Pharmaceutical Development Company, Inc., which licenses a nanoparticle technology that is described in patent US-20150190485. All other authors declare they have no competing interests.

## Data availability

Count matrices, code, and code outputs for both bulk RNA sequencing datasets is available at: https://github.com/shea-lab/Viola_2026_NP.Fibrosis. All other data available upon reasonable request to the authors. Raw sequencing data will be available on Omnibus pending acceptance.

## Notes

https://github.com/shea-lab/Viola_2026_NP.Fibrosis

