## Supplementary material for "“Immunomodulatory nanoparticles elicit antifibrotic monocyte activation to resolve murine pulmonary fibrosis”": SI Figures, Tables, Methods

**SUPPLEMENTAL FIGURES**

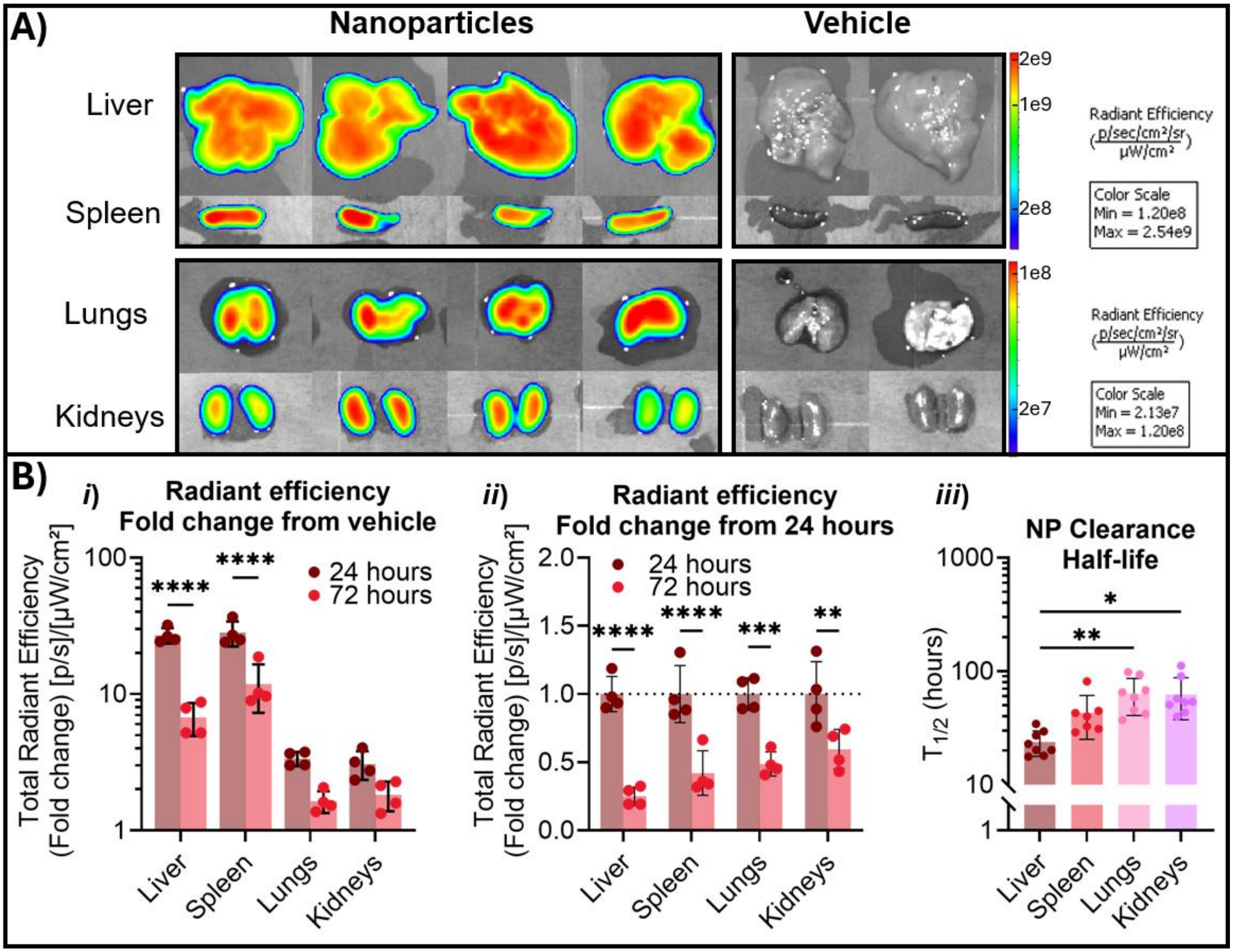


**Figure S1. NPs degrade within days from solid organs.** WT male 6-8 week C57BL/6J mice received 1 dose of fluorescently conjugated (Cy5.5) NPs followed by whole-organ fluorescence imaging 24 or 72 hours post-NPs to determine NP biodistribution and degradation rates. NPs accumulated in blood-filtering organs as expected with predominant distribution to the liver and spleen (**Figure S2A**). NPs showed degradation from these sites within 72 hours with calculated half lives of 23.74 ± 5.99 (SD) hours in the liver, 43.06 ± 17.95 (SD) hours in the spleen, 62.48 ± 25.15 (SD) hours in the kidneys, 63.46 ± 22.78 (SD) hours in the lungs.

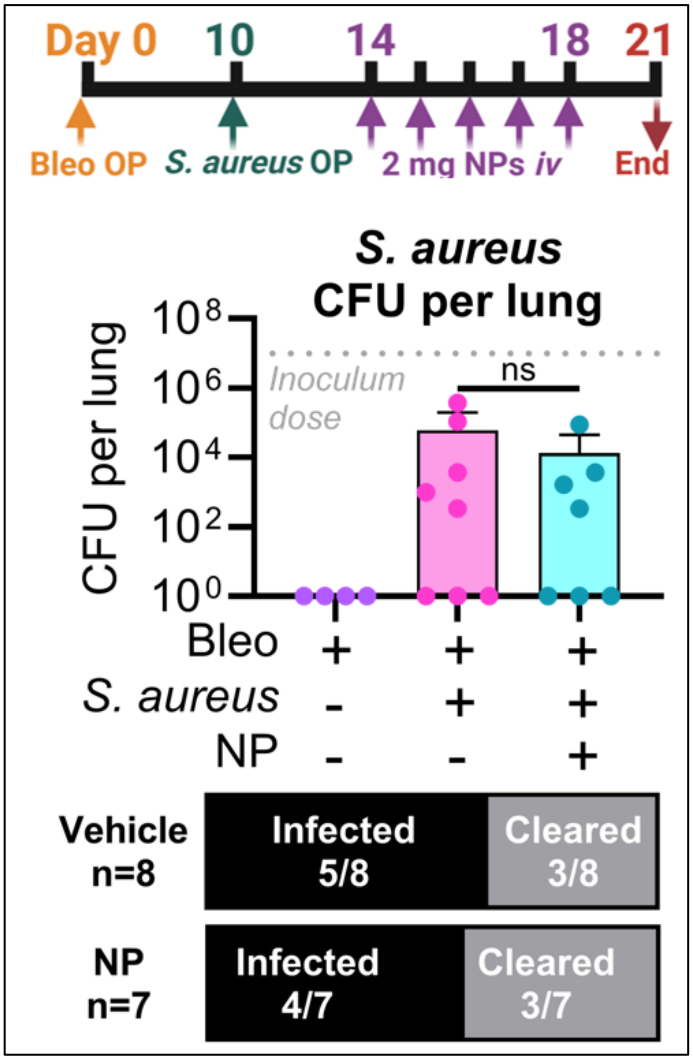


**Figure S2. NPs do not increase suceptibility to *S. aureus* in fibrotic animals.** WT male 6-8 week C57BL/6J mice were inoculated with S. aureus oropharyngeally (ST8:USA300, $10^7$ colony forming units (CFU)/mouse) on day 10 post-bleomycin. NPs were delivered *i.v.* on days 14-18 and CFU was counted on Day 21 from whole lung homogenate. NPs did not affect *S. aureus* clearance rates or total lung

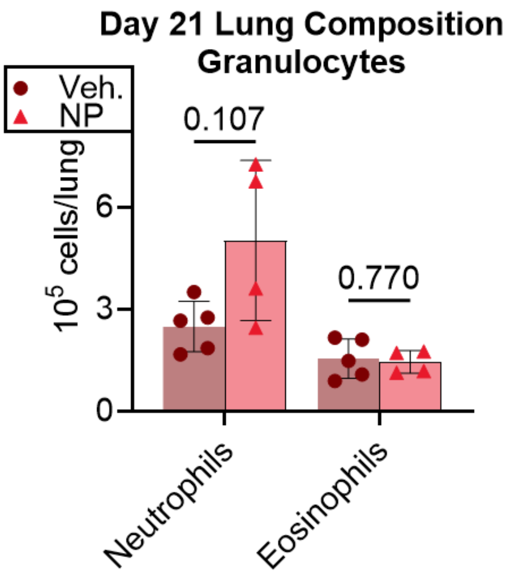


**Figure S3.** Granulocyte numbers were not significantly increased with NP treatment.

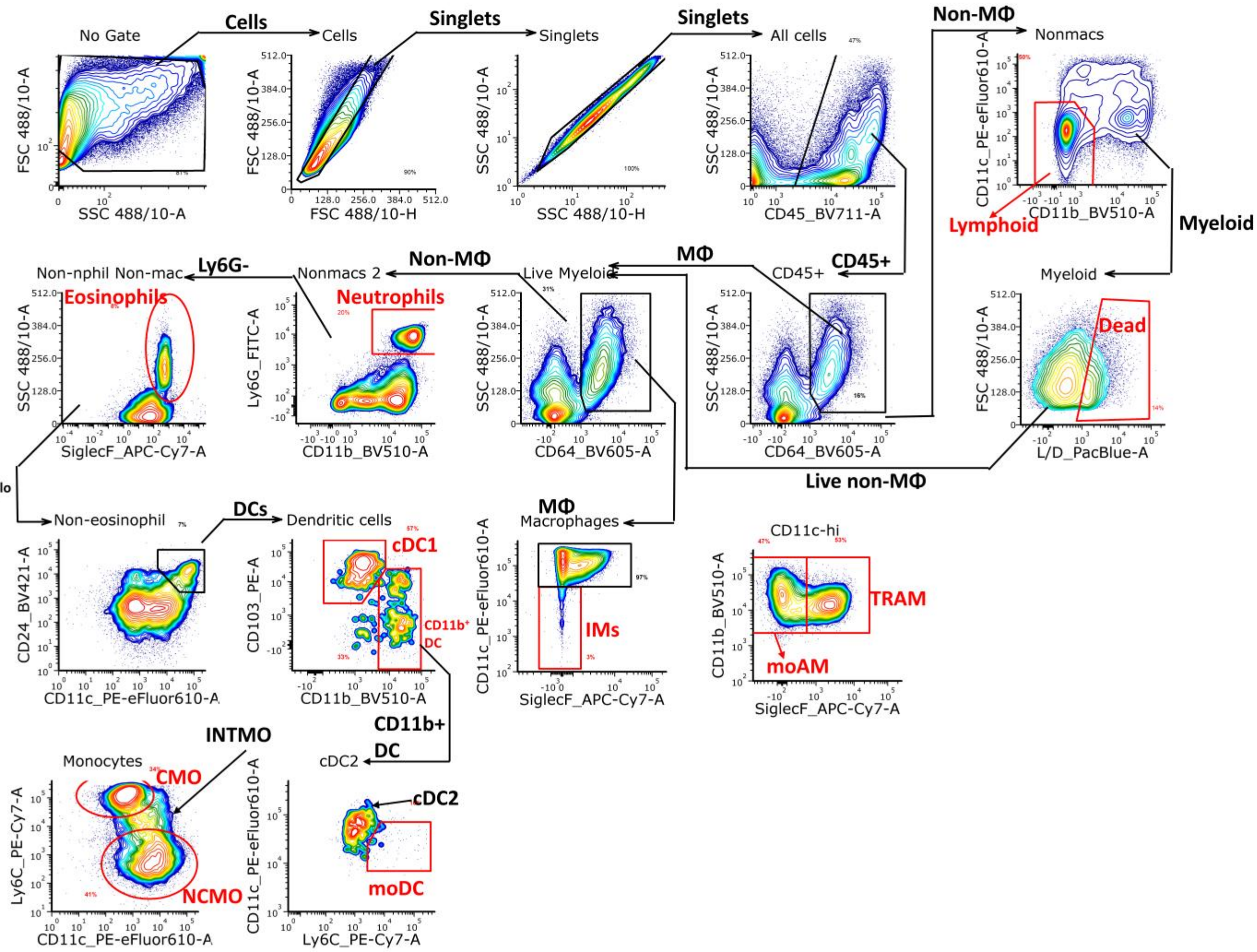


**Figure S4.** Gating strategy for whole lung flow cytometry assay (1/2).

| | |
|---|---|
| MΦ | Macrophage |
| DC | Dendritic cell |
| cDC1 | Conventional dendritic cell, type I |
| cDC2 | Conventional dendritic cell, type II |
| moDC | Monocyte-derived dendritic cell (equivalent to Ly6C++ DC) |
| IM | Interstitial macrophage |
| TRAM | Tissue-resident alveolar macrophage |
| moAM | Monocyte-derived alveolar macrophage |
| CMO | Classical monocyte |
| INTMO | Intermediate monocyte |
| NCMO | Nonclassical monocyte |

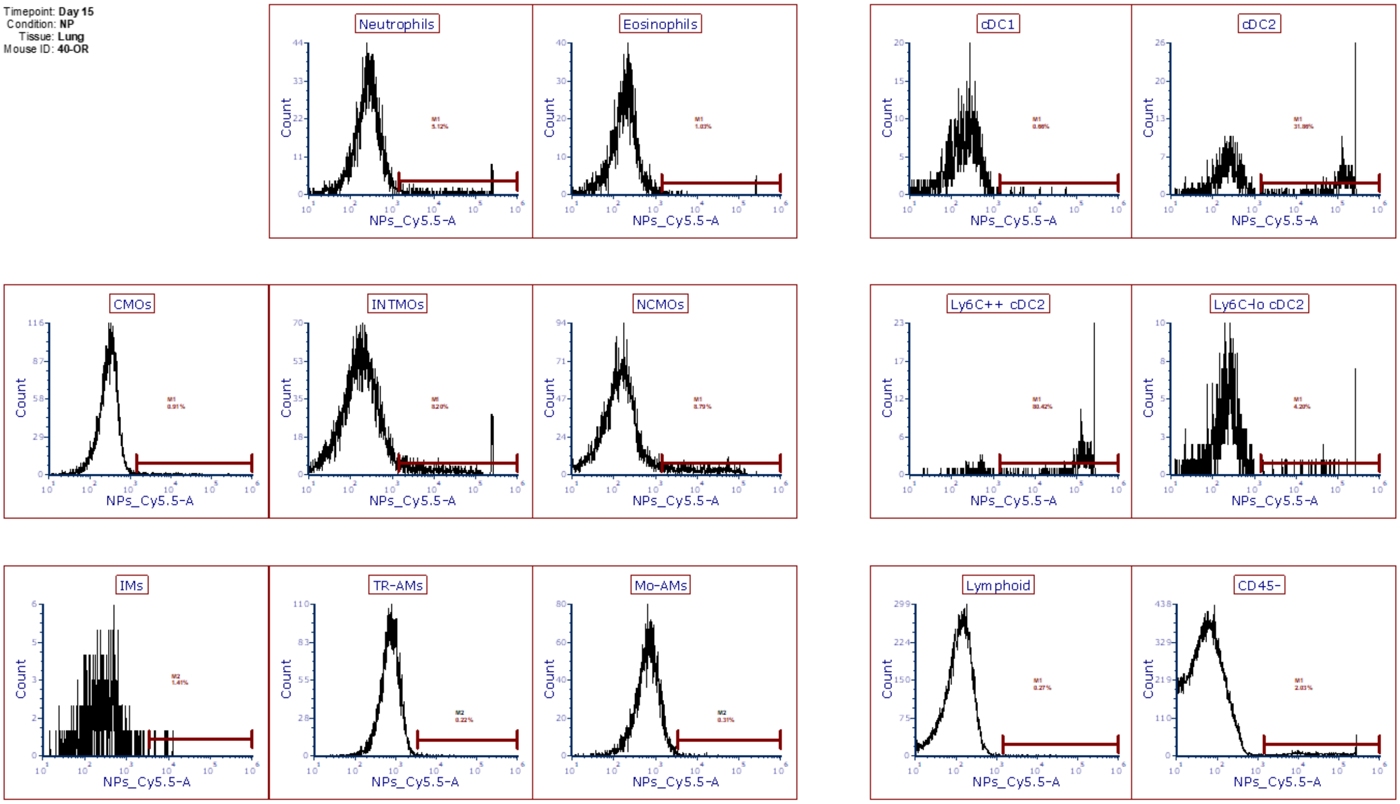


**Figure S5.** Gating strategy for whole lung flow cytometry assay (2/2).

| | |
|---|---|
| MΦ | Macrophage |
| DC | Dendritic cell |
| cDC1 | Conventional dendritic cell, type I |
| cDC2 | Conventional dendritic cell, type II (equivalent to Ly6Clo DC) |
| moDC | Monocyte-derived dendritic cell (equivalent to Ly6C++ DC) |
| IM | Interstitial macrophage |
| TRAM | Tissue-resident alveolar macrophage |
| moAM | Monocyte-derived alveolar macrophage |
| CMO | Classical monocyte |
| INTMO | Intermediate monocyte |
| NCMO | Nonclassical monocyte |

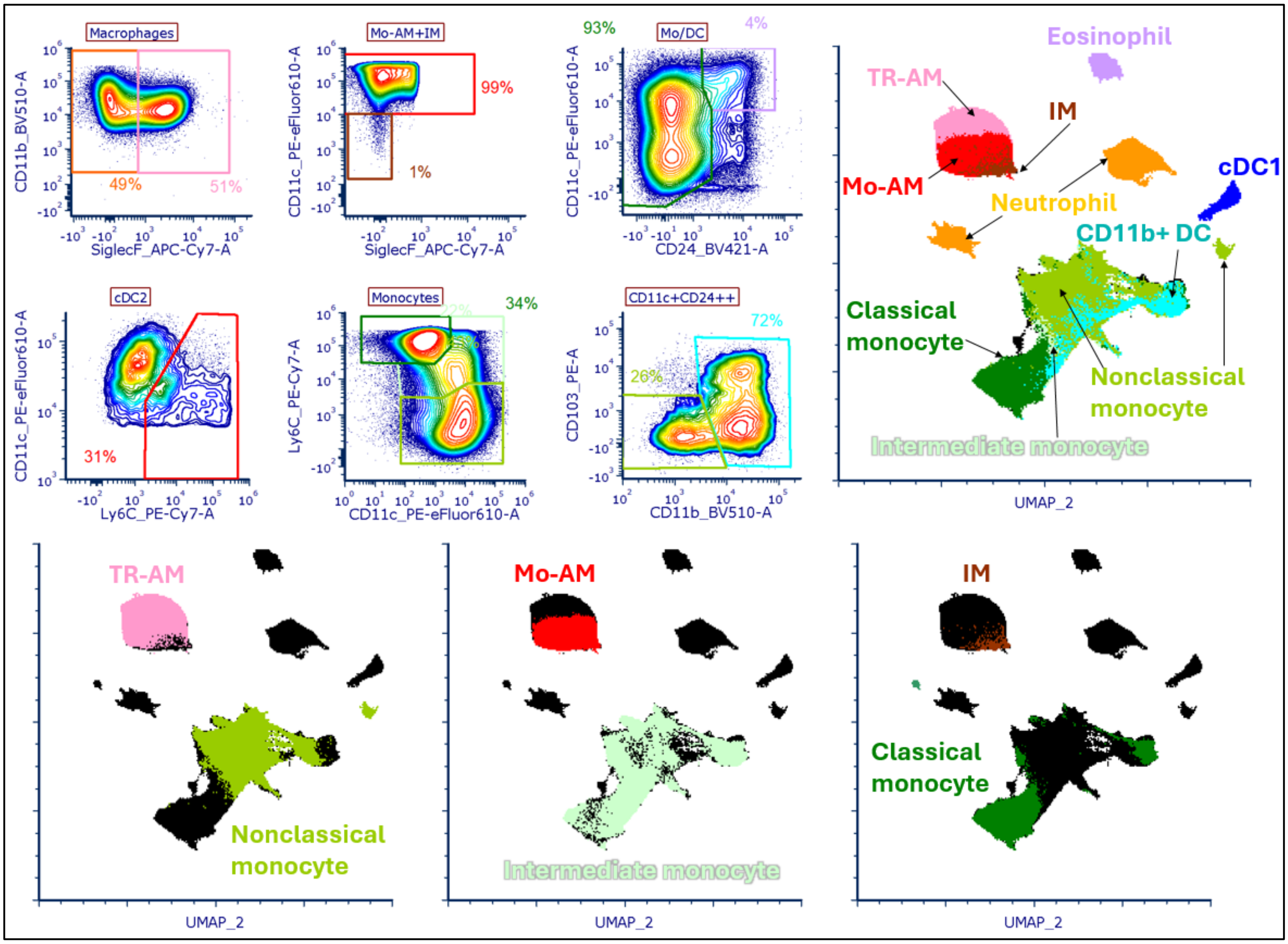


**Figure S6.** Gating strategy for Uniform Manifold Approximation Projection (UMAP) plot of lung flow cytometry data.

### SUPPLEMENTAL TABLES

**Table S1. Genes regulated by NPs in M2-polarized MΦs**

| Function | Increased by NPs | Decreased by NPs | References |
|---|---|---|---|
| Matrix components | *Col20a1, Cspg4, Col4abp3* | *Col14a1, Col18a1, Col27a1, Col4a5, Fn1, Chsy1, Cspg5, Dag1* | |
| Matrix degradation | *Mmp12, Hyal1, Hyal2, Ctsk, Mfge8* | | [1–5] |
| Immunomodulatory mediators | *Ahr, Arnt, Tiparp, Il1rn, Ptges2, Smurf1, Tollip, Ldha, Cav1, Cd36, Igfbp2, Cd22, Cd276* | | [6–13,13–22] |
| Profibrotic mediators | | *Tgfb1, Tgfb3, Postn, Ccl12, Cxcl1, Cxcl2, Sting, Gas6, Mgl2, Cd44, Ccr2, Egfr, Fgf2, Mmp8, C1qa, C1qb, C1qc* | [24–32], [23,33–38] |
| M2 phenotype | | *Arg1, Retnla, Mrc1, Mertk, Cd163, Il6ra, Csf1r, Igf1* | [33,34,39] |
| M1 phenotype | *Il1b, Stat1, Jak2* | | [40] |
| Transcription factors | *Pparg, Stat3, Cebpg, Cebpa, Jag1, Jak1* | Stat1, Stat2, Irf3, Irf4, Jag2, Jak2, Jak3 | |
| Histone editing enzymes | *Kdm1a, Kdm3a, Kdm5a, Kdm5b, Kdm5c, Kdm6b, Kat5, Kat6b, Kat7, Kat8, Dnmt3a, Dnmt3aos* | Hdac1, Hdac5, Hdac6, Hdac9, Hdac10, Kdm2b, Kdm3b, Kat14, Kat2b, Dnmt1 | |

**Table S2. KEGG Pathways regulated by NPs in M2-polarized MΦs.**

| | Increased with NPs | Decreased with NPs |
|---|---|---|
| **Metabolism** | "Glycolysis / Glyconeogenesis"[42]<br>"Pyruvate metabolism"[43]<br>"Amino sugar and nucleotide sugar metabolism"[44]<br>"Biosynthesis of nucleotide sugars" | "Arginine and proline metabolism"[45]<br>"Fatty acid metabolism"[45,46]<br>"Inositol phosphate metabolism"[45]<br>"N-glycan biosynthesis" |
| **Signaling** | "HIF-1 signaling pathway"[47]<br>"mTOR signaling pathway"[47]<br>"NF-κB binding"<br>"p53 signaling pathway"[48]<br>"MAP kinase phosphatase activity" | "PI3K-Akt signaling pathway"[45,49]<br>"MAPK signaling pathway"[45,50]<br>"VEGF signaling pathway"<br>"Toll-like receptor signaling pathway" |

***Table S3. Genes regulated by NPs in murine lungs.***

| Cluster | Differentially expressed genes |
|---|---|
| **Cluster 1**:<br>Decreased over time in NP-treated lungs | **ECM degradation**<br>*Hyal1, Mmp12, Mmp13, Mmp25, Lair1*<br><br>**Inflammation resolution**<br>*Alox12, Cybb, Ccdc88a, Acod1, Il10ra, Il10rb, Nrros, Tnfaip3* (AP1), *Tnfrsf1b, Socs3*, *Il7r*, *Nos3*, *Ptger4*, *Plaa* |
| **Cluster 3**:<br>Increased over time in vehicle-treated lungs | **ECM Components**<br>Col1a2, Col3a1, *Col5a3*, *Col6a3*, *Col12a1*, *Col14a1*, *Col16a1*, *Col28a1*, *Lama2, Lamb1, Lamb2, Matn4, Mgp, Ltbp3, Adamts2*<br><br>**M2 macrophage phenotype**<br>*Mgl2*, *Retnla*, *Loxl2*, *Cd209b* (analogue of human DC-SIGN) |

***Table S4. Highlighted enriched pathways in murine NP-treated lungs by temporal trajectory cluster***

| **Cluster** | **Enriched Pathways** |
|---|---|
| **Cluster 1**:<br>Up in NP-treated lungs | **KEGG Pathways**<br>"Chemokine signaling pathway"<br>"Leukocyte transendothelial migration"<br>"Fc gamma R-mediated phagocytosis"<br>"Regulation of actin cytoskeleton"<br>"NOD-like receptor signaling pathway"<br>"NF-kappa B signaling pathway"<br>"Cytokine-cytokine receptor signaling pathway"<br>"Hematopoietic cell lineage"<br>"Focal adhesion"<br>"Phagosome"<br>"JAK-STAT signaling pathway"<br>"Notch signaling pathway"<br>"Efferocytosis"<br>"Sphingolipid signaling pathway"<br>"C-type lectin receptor signaling pathway"<br><br>**GO Pathways**<br>"collagen binding"<br>"nuclear retinoic acid receptor binding"<br>"vascular endothelial growth factor receptor binding"<br>"peroxisome proliferator activated receptor binding"<br>"phospholipase activator activity"<br>"histone modifying activity"<br>"phosphoprotein binding"<br>"phospholipase binding"<br>"Toll-like receptor binding"<br>"CXCR chemokine receptor binding"<br>"integrin binding"<br>"histone H3K9 demethylase activity"<br>"glycolipid binding" |
| **Cluster 2**:<br>preserved in NP- but not vehicle-treated lungs from Day 14 to Day 21 | **KEGG Pathways**<br>"Tnf signaling pathway"<br>"Adipocytokine signaling pathway"<br>"Viral protein interaction with cytokine and cytokine receptor"<br><br>**GO Pathways**<br>"antioxidant activity"<br>"growth factor receptor binding" |
| **Cluster 3**:<br>Up in vehicle-treated lungs | **KEGG Pathways**<br>"Cardiac muscle contraction"<br>"Chemical carcinogenesis – reactive oxygen species"<br>"Cytoskeleton in muscle cells"<br>"ECM-receptor interaction"<br>"Oxidative phosphorylation"<br>"Protein digestion and absorption"<br>"Proteasome"<br><br>**GO Pathways**<br>"carbohydrate binding"<br>"extracellular matrix structural constituent conferring tensile strength" |

| | |
|---|---|
| | “glycosaminoglycan binding”<br>“heparin binding”<br>“platelet-derived growth factor binding”<br>“proton-transporting ATPase activity, rotational mechanism”<br>“sulfur compound binding” |
| **Cluster 4**: preserved in vehicle- but not NP-treated lungs from Day 14 to Day 21 | **KEGG Pathways**<br>No KEGG pathways were enriched.<br><br>**GO Pathways**<br>“Calmodulin binding” |

### DETAILED MATERIALS AND METHODS

***qRT-PCR.*** RNA was isolated using Trizol (Invitrogen) according to manufacturer's instructions. RNA quantity and purity were assessed on a nanodrop spectrophotometer (Thermo). Transcript expression was analyzed *via* reverse transcriptase quantitative real-time PCR using Lunaprobe reagent (New England Biolabs) on a Quantstudio3 thermocycler (ABI). Expression analysis was conducted using a ΔΔct calculation and shown as fold-change from NP-treated vs. PBS-treated cells from the same donor animal. Transcripts were normalized to expression level of the housekeeping gene RPL38. Primer sequences are specified in Table S5.

**Table S5. PCR Primer sequences**

| | |
|---|---|
| *Rpl38* | Integrated DNA technologies, Hs.PT.58.40595235.gs |
| *Fizz1* | Forward - 5'-TCCAGCTAACTATCCCTCCACTGT-3'<br>Reverse - 5'-AGCCACAAGCACACCCAGTAG-3'<br>Probe - 5′-ATGAACAGATGGGCCTCCTGCCCT-3' |
| *Il6* | Integrated DNA technologies, Mm.PT.58.10005566 |
| *Cd206* | Forward - 5'-AGTGGCTTTGGTTGAACGAC-3'<br>Reverse - 5'-CCAAAGGCCCGAAGATGAAG-3'<br>Probe - 5'-CCGTTCACCAGACCCATCGCCT-3' |
| *Tnfa* | Integrated DNA technologies, Mm.PT.58.12575861 |
| *Ccl3* | Forward - 5'-CTTCTCCTACAGCCGGAAGATTC-3'<br>Reverse - 5'-GCCGGTTTCTCTTAGTCAGGAA-3'<br>Probe- 5'-GAAACCAGCAGCCTTTGCTCC-3' |
| *iNos* | Integrated DNA technologies, Mm.PT.58.43234558 |

**Animal models:** Male wild type C57BL/6J mice and female wild type Balb/c mice 6-8 weeks of age were purchased from Jackson Laboratories (Bar Harbor, ME). Male B6.Cg-Tg(Col1a1*2.3-GFP)1Rowe/J (Col-GFP) mice on C57BL/6J background were bred in-house and used between 6-12 weeks of age. All animal procedures were conducted in compliance with the University of Michigan's Institutional Animal Care & Use Committee (IACUC) guidelines. Animals were housed in specific pathogen free conditions with *ad libitum* access to chow and water. All control and fibrotic animals were provided ad libitum wet diet (DietGel® by ClearH2O) to prevent dehydration due to bleomycin treatment. Bleomycin was administered oropharyngeally under 3-4% isoflurane at 0.0125 U/mouse in 50 uL (0.5 U/kg for a 25 g mouse). NPs were delivered intravenously to anesthetized mice *via* retro-orbital sinus or to awake mice *via* lateral tail vein.

**Nanoparticle fabrication:** NPs were fabricated using an oil-in-water single emulsion technique as previously described[51]. 50:50 Poly(DL-lactide-*co*-glycolide) (PLG) was purchased from Evonik (LACTEL® 50:50 DL-PLGA IV 0.55-0.75 (legacy LACTEL® B6013-2), acid terminated, intrinsic viscosity 0.55-0.75 dL/g in hexafluoroisopropanol). PLG was dissolved in dichloromethane (CAS# 75-09-2). The organic phase was mixed with an aqueous solution of 2% w/v poly(ethylene-alt-maleic anhydride) (PEMA) with MW ~400 kDa (CAS# 9006-26-2, Polysciences Inc. #02308). An emulsion was created *via* probe sonication for 30 seconds at 100% amplitude (Cole-Parmer CPX130 Ultrasonic Processor with a Cole-Parmer mm probe with stepped tip) and the emulsion was then mixed into a 200 mL aqueous solution of 0.5% w/v PEMA. Solvent was allowed to evaporate while mixing at room temperature for 16 hours. NPs were then washed 4 times in 0.1M sodium bicarbonate (Polysciences NC1489512), resuspended in water plus 25% v/v cryoprotectant (cryoprotectant: 15% w/v D-mannitol and 20% w/v sucrose in milliQ water), frozen at -80 °C, lyophilized for 48 hours, and stored at room temperature under vacuum protected from light until use. For intravenous injections, NPs were washed in 1 mQ sterile $H_2O$ to remove cryoprotectant and resuspended in sterile Dulbecco's Phosphate Buffered Saline (DPBS) (Fisher Scientific 14-190-250) at 10 mg/mL. NPs were filtered through a flow cap tube (70 µm) immediately prior to *i.v.* delivery. NP size and zeta potential were measured with a Zetasizer (Malvern Panalytical) according to manufacturer instructions. For fluorescence detection of NPs, Cy5.5-NHS, a small molecule dye with a reactive linker, was purchased from Lumiprobe and conjuaged to acid-terminated PLG using EDC-NHS chemistry (EDC, 1-Ethyl-3-(3-dimethylaminopropyl)carbodiimide, CAS# 25952-53-8; NHS, N-Hydroxysuccinimide sodium salt, CAS# 106627-54-7) as we have described elsewhere[52]. 20% w/w Cy5.5-conjugated PLG was used to fabricate fluorescent NPs.

**Bone marrow derived macrophages (Figure 1C):** Bone marrow (BM) was harvested from tibias and fibulas of male B6.Cg-Tg(Col1a1*2.3-GFP)1Rowe/J (Col-GFP) mice on C57BL/6J background. BM was pushed out of cutoff long bones using an 18G needle and then lysed, filtered (100 µm), counted, and plated in non-tissue culture treated 10 cm diameter petri dishes at 15 million cells in 10 mL BMDM media per plate. BMDM media consisted of, by volume, 30% L929 fibroblast conditioned media, 59% IMDM, 10% fetal bovine serum, and 1% penicillin/streptomycin. After seeding, BMDMs were allowed to differentiate in BMDM media for 6 days in a cell culture incubator (37 °C, 95% humidity, 5% $CO_2$). On day 6, NPs (200 µg/mL in 20 mLs for total 8 mg per dish) or vehicle were added to the dish and incubated for 24 hours. Finally, BMDM supernatant was collected, spun to remove debris and particles, and used for the scratch wound assay. BMDMs were lysed with TRIzol® reagent (Zymo Research) for qRT-PCR.

**Fibroblast scratch wound assays:** Fibroblasts were cultured from the same donors as the bone marrow, i.e., male Col-GFP mice 6-12 weeks of age. Fibroblasts were expanded from mouse lungs using a "crawl-out culture" method as previously described[53]. Briefly, lung lobes were finely minced to <1 mm and cultured in a tissue culture treated flask with RPMI-1640 with 10% v/v FBS and 1X penicillin/streptomycin. Once tissue pieces were firmly attached to the flask after 24-48 hours, cells were rinsed to remove debris and red blood cells. Media was changed every 2-3 days for 14 days. Then, fibroblasts were passaged with trypsin and plated at 62,500 cells/$cm^2$ in Incucyte® Imagelock 96-well Microplate (Sartorius BA-04856) until reaching confluence after 16 hours. When cells reached confluence, scratch wounds were created using the Incucyte® WoundMaker (Sartorius) according to manufacturer instructions. Fibroblasts were washed to remove debris and NP- or vehicle-treated BMDM conditioned media was added to each well with 6 technical replicates per animal per condition. Two fluorescence and brightfield images per well were collected every 3 hours for 48 hours using the Incucyte® Live-Cell Analysis System. Scratch wound confluence and fibroblast area were quantified using the Incucyte® Scratch Wound Analysis Software Module (Sartorius, Cat. No. 9600-0012).

**Bone marrow derived macrophages (Figure 1D-F):** Bone marrow (BM) was harvested from tibias and fibulas of female BALB/c mice 6-12 weeks of age. BM was pushed out of cutoff long bones using an 23G needle, then lysed, filtered (70 µm), counted, and plates on non-tissue culture treated 10 cm dish at 2 million cells in 10 mL macrophage differentiation media (LCM: RPMI 1640 supplemented with 10 v/v% FBS, 1x P/S, and 4 mM L-glutamine; MPM: 80 v/v% LCM supplemented with 20 v/v% L929 fibroblast conditioned media) and were cultured for 8 days. Macrophages were polarized for 24 hours on day 8 to M2 with stimulation by 10 ng/mL each of IL-4 and IL-13 (ThermoFisher Scientific). On day 9, M2 macrophages were treated with 200 ug/mL NPs for 24 hours followed by rinsing to remove residual NPs and RNA extraction using the Direct-Zol™ RNA MiniPrep Plus with Zymo-Spin™ IIICG Columns (Zymo Research Corporation R2070) according to manufacturer instructions. RNA sequencing was performed as described above.

**Collagen quantitation:** Whole lung lobes were excised from large airway and trachea, and all 5 lobes were degraded for 20 hours at 95 °C in 1 mL 6M hydrochloric acid (CAS# 7647-01-0) using an aluminum beads (Lab Armor) in a dry block incubator. Whole lung collagen content was assessed from this hydrolysate using the Quickzyme Total Collagen Assay (Ilex Biosciences QZBTOTCOL1) according to manufacturer instructions.

**µCT ventilation perfusion imaging:** Murine µCT was performed on day 29 post-bleomycin in NP or vehicle-treated animals to assess lung perfusion. Male BL6 wilde type animals aged 6-8 weeks were anesthetized with 100 mg/kg ketamine, intubated, and transferred to a phantom-calibrated µCT under 1.5% isoflurane. 2 minute lung geometry scans are performed and the mouse is administered reversal agents. Finally, a phantom geometry calibration is taken for delivery to 4DMedical (Australia). 4DMedical applies a proprietry algorithm to quantify air velocity and lung perfusion in calibrated µCT images. These analysis files, including all analyzed images from coronal and axial CT, are included in Supplemental files.

**Lung Bulk RNA sequencing:** Whole lung lobes were excised from large airway and trachea, and all 5 lobes were homogenized at 15000 rpm for 1-3 minutes in 1 mL TRIzol® reagent (Zymo Research) to achieve a pipetteable solution. This solution was spun to remove solids and 150 µL supernatant was diluted with 650 µL TRIzol®. Then, RNA was extracted using the Direct-Zol™ RNA MiniPrep Plus with Zymo-Spin™ IIICG Columns (Zymo Research Corporation R2070) according to manufacturer instructions.

RNA sequencing (lungs and macrophages): RNA quality and concentration were quantified by the University of Michigan Advanced Genomics Core (AGC) *via* RNA gel (Agilent TapeStation). AGC conducted library prep with poly-A enrichment and next-generation sequencing (Illumina NovaSeq) according to manufacturer instructions. For lungs, 530 million reads were generated across 12 samples for a mean sequencing depth of 44 ± 7.7 million reads/sample. ***Data analysis.*** AGC aligned sequences to generate raw count matrices. Raw counts were processed in RStudio using DESeq2 (https://github.com/thelovelab/DESeq2). First, a DESeq results object was created using genes with at least 5 counts per sample. To find differentially expressed genes (DEGs), we used DESeq2's Likelihood Ratio Test with independent hypothesis weighting (https://github.com/nignatiadis/IHW). We identified clusters of differentially expressed genes with DEGreport (https://github.com/lpantano/DEGreport). Gene set enrichment analysis was performed using ClusterProfiler (https://guangchuangyu.github.io/software/clusterProfiler/) and heatmaps were generated using ComplexHeatmap (https://github.com/jokergoo/ComplexHeatmap). Code and count matrices for both datasets are available at https://github.com/shea-lab/Viola_2026---NPs_in_Fibrosis.

**IVIS Imaging:** Whole organs were excised from sacrificed animals and fluorescence images were taken using an IVIS™ system with LivingImage software using Low brightness, excitation 640 nm/emission 700 nm, 0.5 s exposure time for all images. Regions of interest were outlined manually to extract radiant efficiency for each organ.

***S. aureus* infection model:** NP studies were conducted identically to those described prior, with the addition of oropharyngeal *S. aureus* inoculation (strain USA300, 1e5 CFU) on day 10 post-bleomycin. On day 21, S. aureus was quantified from whole homogenized lung lobes via colony forming assay as described previously[54]. Homogenized lung tissue was serially diluted to 8 orders of magnitude and plated on Nutrient Agar (BD Difco) for 24 hours at 37 °C. Then, *S. aureus* colony forming units were counted manually with an inverted brightfield microscope.

**Flow cytometry**

**Table S6. Flow cytometry antibodies**

| Target | Color | Ab clone | Ab isotype | Vendor | Cat# |
|---|---|---|---|---|---|
| CD24 | BV421 | M1/69 | Rat IgG2b, κ | BioLegend | 101825 |
| CD11b | BV510 | M1/70 | Rat IgG2b, κ | BioLegend | 101245 |
| CD64 | BV605 | X54-5/7.1 | Mouse IgG1, κ | BioLegend | 139323 |
| CD45 | BV711 | 30-F11 | Rat IgG2b, κ | BioLegend | 103147 |
| Ly6G | FITC | 1A8 | Rat (LEW) IgG2a, κ | BD Biosciences | 551460 |
| CD103 | PE | 2E7 | Armenian hamster / IgG | ThermoFisher | 12-1031-82 |
| CD11c | PE/eFluor610 | N418 | Armenian hamster / IgG | ThermoFisher | 61-0114-82 |
| Ly6C | PE/Cy7 | HK1.4 | Rat IgG2c, κ | BioLegend | 128018 |
| SiglecF | APC-Cy7 | E50-2440 | Rat LOU, also called Louvain, LOU/C, LOU/M IgG2a, κ | BD Biosciences | 565527 |

***Lung processing.*** Lung lobes were minced on ice into pieces <1 mm and digested in 50 mL conical vials with 15 mL digest solution for 40 minutes at 37 °C with moderate agitation. Digest solution consisted of Roswell Park Memorial Institute (RPMI)-1640 medium with L-glutamine (Fisher Scientific 61870127) plus 1 mg/mL collagenase A (Sigma Aldrich 10103578001) and 17 kU/mL deoxyribonuclease I from bovine pancreas (Sigma Aldrich D4263-1VL). Digested lungs were diluted in RPMI, dissociated with a 16 G needle, pushed through a 100 µm filter, pelleted and washed with RPMI, and resuspended in 3 mL ACK lysis buffer (ThermoFisher A1049201) for 5 minutes at room temperate to remove red blood cells. After lysis, cells were washed with RPMI, filtered through 100 µm cell strainer, and counted. 1 million live cells per lung were aliquotted onto a 96-well round bottom microplate for antibody staining.

***Spleen processing.*** Spleens were mashed through a 100 µm cell strainer, washed, and lysed with 10 mL ACK lysis buffer for 5 minutes at room temperature. Cells were washed, filtered (100 µm), counted, and plated at 1 million live cells/sample.

***Blood processing.*** Peripheral blood was collected into EDTA microvettes (Kent Scientific MTSC-EDTA) *via* terminal cardiac puncture on anesthetized animals (5% isoflurane) followed immediately by euthanasia with cervical dislocation. Blood was lysed in 5 mL ACK lysis buffer (5 minutes at room temperature), washed twice, filtered (100 µm), counted, and plated at 1e6 live cells/sample.

***Antibody staining.*** Cells were resuspended in live/dead cocktail and incubated protected from light for 10 minutes at 4 °C (live/dead cocktail: 1:100 Viobility™ 405/452 Fixable dye, Miltenyi Biotec 130-130-420 in sterile DPBS with 0.5 M Ethylenediaminetetraacetic acid, (EDTA)). Cells were washed with staining buffer (DPBS + 0.5M EDTA + 0.5% w/v sterile bovine serum albumin) and incubated in 50 µL staining buffer + 1:100 TruStain FcX™ (anti-mouse CD16/32) Antibody (BioLegend 101319) for 10 minutes on ice followed by direct addition of 50 µL 2x antibody cocktail and a further 30 minutes incubation at 4 °C protected from light (2x antibody cocktail: BD Horizon™ Brilliant Stain Buffer (BD Biosciences 563794) + 1:100 FcX™ + all antibodies listed below in Table S7). After antibody staining, cells were washed twice, fixed with 4% PFA on ice for ten minutes, washed, and stored at 4 °C protected from light until data collection within 72 hours. ***Data collection and analysis.*** All single stained compensation controls and samples were collected during one session on a ZE5 Cell Analyzer (Bio-Rad). Compensation and data analysis were performed in FCS Express™ (De Novo Software by Dotmatics). Gating strategies are shown in Supplemental Figures S4-S6.

***Table S7.*** *Lung celltype markers*

| | BV711 | SSC-488-A | BV605 | PE-eFl.610 | BV421 | BV510 | PE | PE/Cy7 | FITC | APC/Cy7 |
|---|---|---|---|---|---|---|---|---|---|---|
| Celltype | **CD45** | **SSC-A** | **CD64** | **CD11c** | **CD24** | **CD11b** | **CD103** | **Ly6C** | **Ly6G** | **SiglecF** |
| Eosinophil | + | ++ | - | | + | | | | - | +++ |
| Neutrophil | + | ++ | - | | | +++ | | | +++ | |
| TR-AM | + | +++ | + | +++ | | lo/- | | | | +++ |
| Mo-AM | + | +++ | + | +++ | | + | | | | lo/- |
| IM | + | +++ | + | + | | + | | | | - |
| CMO | + | lo | - | lo/- | lo/- | | | +++ | - | |
| INTMO | + | lo | - | + | lo/- | | | + | - | |
| NCMO | + | lo | - | +++ | lo/- | | | lo/- | - | |
| cDC1 | + | lo | - | +++ | +++ | - | +++ | | - | |
| cDC2 | + | lo | - | +++ | +++ | +++ | lo/- | | - | |

**Statistics:** All statistical calculations were performed in GraphPad Prism 10.4.0. Statistical tests for specific datasets are specified in figure captions. For all figures, *p≤0.05, **p≤0.01, ***p≤0.001, ****p≤0.0001.

**Code:** Count matrices, code, and code outputs for both bulk RNA sequencing datasets is available at: https://github.com/shea-lab/Viola_2026_NP.Fibrosis.

54. Warheit-Niemi, H. I. *et al.* Fibrotic lung disease inhibits immune responses to staphylococcal pneumonia via impaired neutrophil and macrophage function. *JCI Insight* **7**, e152690.
